# Molecular dissection of the cellular response to mechanical stress

**DOI:** 10.1101/539262

**Authors:** Noam Zuela-Sopilniak, Jehudith Dorfman, Yosef Gruenbaum

## Abstract

The response of cells to mechanical stress is crucial for many cellular functions, yet its molecular mechanisms are not yet fully understood. Previous studies of the cellular response to mechanical stress were performed on cultured cells or isolated muscle fibers devoid of cell and/or tissue contexts. Thus, the emerging results were limited to the specific cell types or tissues analyzed and dependent on the growth matrix elasticity. In the present study, we looked for changes in early gene expression in response to mechanical whole body stretching of living *C. elegans*. Our transcriptome analysis revealed upregulation of genes involved in cuticle development, stress response and several signaling pathways such as WNT, TGFβ, AMPK and Hedgehog signaling. These findings indicate that protecting against mechanical insults entails providing additional support to the mechanically resilient protective cuticle, and that proper recovery from mechanical stretching requires an intact Hedgehog signaling pathway. Recent findings suggest an important role for the nuclear lamina in mediating cellular mechanical response. The nuclear lamina is composed mainly of lamins, which are nuclear intermediate filament-type proteins needed, among other functions to maintain nuclear integrity. One particular area of interest are laminopathies, which are caused by mutations in lamin. Stretched animals expressing the Emery Dreifuss Muscular Dystrophy (EDMD) linked L535P lamin mutation, showed further upregulation of cytoskeleton organization, cellular respiration and mitochondrial protein-unfolding stress response genes, most likely to compensate for aberrant muscle tissue function. These findings provide a broad multi-dimensional picture of the *in vivo* genetic response of live animals to mechanical stress, highlighting previously unreported mechano-sensitive genes and molecular pathways.

## Introduction

Gene expression in response to mechanical stress was demonstrated on tissue cultured cells including human fibroblasts and rat Müller cells [1,2]. The fibroblasts exposed to mechanical stress activated the MAPK proteins ERK, p38 and JNK and transcription factors such as *AP-1, c-fos* and *c-Jun*, which are involved with osteoblastic differentiation. Gene expression profiling of mechanically stressed Müller cells demonstrated upregulation of the proliferation genes *Areg* and *Atf3*, angiogenic factors such as *Epha2* and Nrn1 and the MAPK pathway. The controlled force resulted in reorganization of the actin cytoskeleton, changes in chromatin compaction and structure, leading to increased transcription. Also affected were processes ranging from protein conformation and assembly, to mechano-sensitive transcription factors (YAP and TAZ) localization to the nucleus [1,2]. In addition to the MAPK pathway, other pathways such as WNT signaling, which can affect proliferation and growth, and TGF-β responded to an external force [3–5].

The nuclear lamina (NL) lies adjacent to the inner nuclear membrane (INM) and is composed of a meshwork of lamins and their associated proteins [6]. Lamins are evolutionarily conserved nuclear intermediate filament (IF) proteins that are involved in nuclear activities ranging from mechanical stability, signal transduction, cell proliferation and chromatin organization to DNA damage repair and transcription [7–9]. Similar to all IF proteins, lamins contain an N-terminal head domain, a central α helical rod domain and a carboxy tail domain containing an immunoglobulin (Ig)-fold, an unstructured region, a nuclear localization signal and a CaaX motif [10,11]. In mammals, 3 genes encode 4 major lamin proteins. The B-type lamins, lamin B1/B2, are encoded by two genes, termed *LMNB1* and *LMNB2* and are expressed in all cells. The A-type lamins, lamins A/C, are encoded by a single gene, termed *LMNA* and are expressed in differentiated cells [12].

There are at least 14 distinct diseases caused by over 600 different missense mutations in the *LMNA* gene. These diseases, collectively termed as laminopathies, are mostly autosomal dominant, and appear postnatally. While originating from a specific *LMNA* mutation, which is globally expressed, many laminopathies show tissue specificity. Laminopathies can be roughly divided into 4 disease categories: the striated muscle diseases, peripheral neuropathies and metabolic diseases affect a specific tissue, while the accelerated aging diseases affect multiple tissues [9,13,14]. The mechanical model of laminopathies tries to explain the muscle-specific degeneration in those diseases [13–15]. This model proposes that the mutant lamin does not support the normal nuclear mechanical response, which leads to nuclear rupture, cell death and tissue deterioration in load-bearing tissues [16]. The multiple lamin-interacting partners, several of which are differentially expressed, may be another factor contributing to tissue specificity of these diseases.

*C. elegans* has one lamin gene (*lmn-1*) encoding a single lamin protein isoform (Ce-lamin), which contains a CaaX motif. Ce-lamin shares many functions with mammalian lamin A such as regulating the mechanical properties of the nucleus, presence in the nucleoplasmic fraction, as well as interacting with binding partners that in mammals interact with lamin A [17,18]. As is the case in humans, dominant expression of specific lamin disease-causing mutations in *C. elegans* leads to phenotypes, which mimic the human corresponding disease phenotypes [19]. A recent study has shown that the mechanical response of the L535P-EDMD-linked lamin mutation is specifically impaired in muscle cells [20] and that the expression of this mutation in relaxed animals causes an overall downregulation of genes involved in muscle organization and function, as well as an impaired pharyngeal structure [21]. Here, we developed a method that allows application of mechanical force to hundreds of living worms. We used it to study the immediate mechano-sensitive transcriptional response to external mechanical strain application, as well as the early transcriptional changes following the recovery process in both WT and L535P animals.

## Results

### Detecting early global mechano-induced transcriptional changes in living *C. elegans*

Applying simultaneous mechanical strain to a large population of living *C. elegans* establishes the mechanically-sensitive transcriptional response at the level of an intact living organism. This method takes into account the effects of neighboring cells and tissues on the response to mechanical strain. In our experimental setup, whole animals are stretched for the duration of 5 minutes. This relatively short time scale ensures that animals are not overly stressed and survived the stretch with no apparent phenotypes. A longer stretching time may lead to mortality accompanied by the initiation of a massive stress response activation related to animal viability. This would mask the specific genetic changes resulting from the application of mechanical stress. Indeed, animal survival following mechanical stretching decreased in correlation with increased stretching time starting from a few hours after stretching occurred (Fig. 1; 97%, 82% and 35% survival following 5, 15 and 30 minutes of mechanical stretching, respectively). Following the 40 minutes recovery period, we were able to detect early mechano-sensitive transcriptional changes alongside the early response of animals to the elimination of strain.

**Figure 1:**
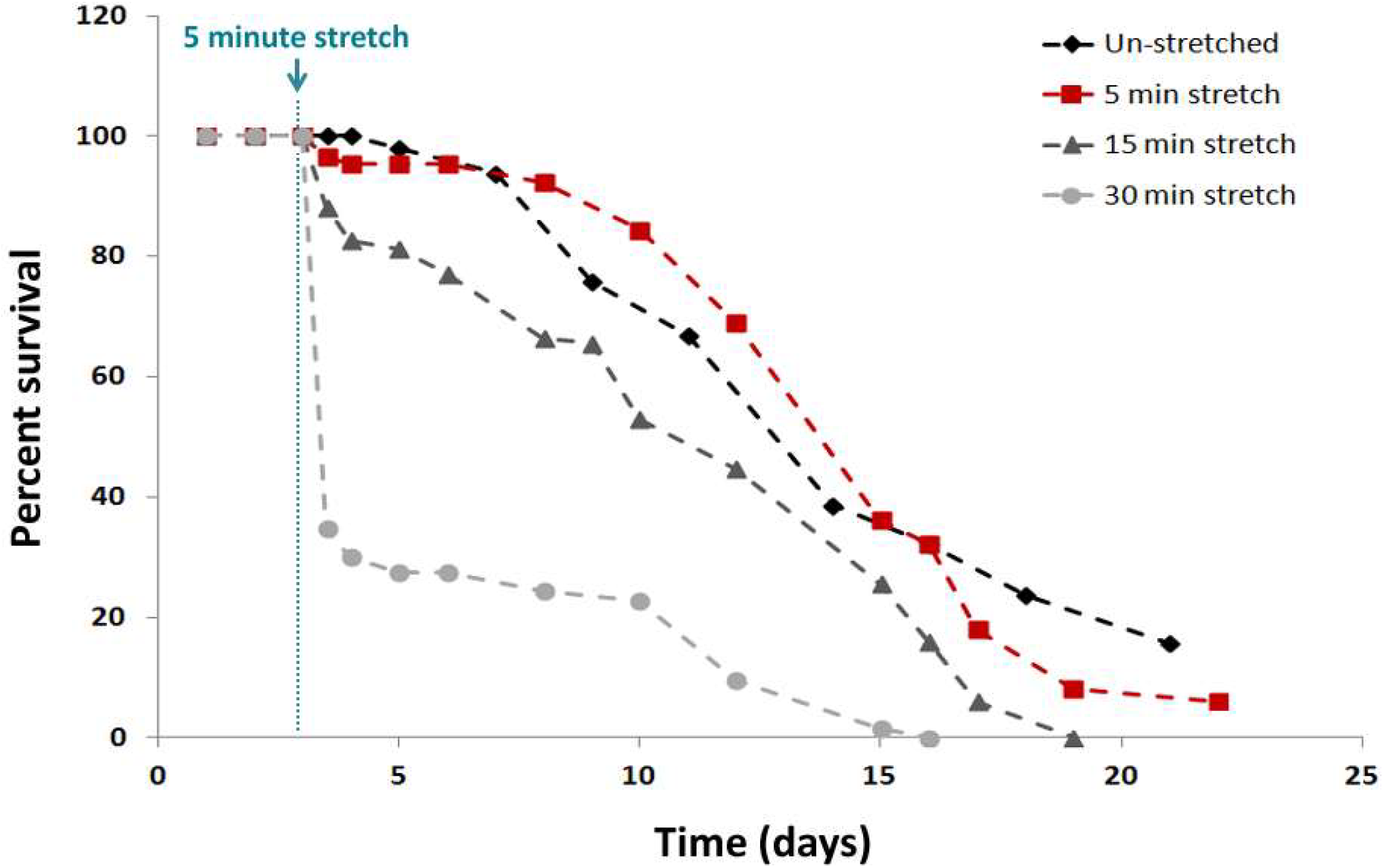
Increasing stretching time decreases animal survival. Survival curves for WT worms at day 2 of development (L4) before and after 40% stretching for 5 (red, n=87), 15 (dark grey, n=144) and 30 (light grey, n=170) minutes. Un-stretched N2 worms (black, n=150) served as controls. Within few hours of stretching, roughly 3%, 18% and 65% lethality was detected in N2 animals exposed to 5, 15 and 30 minutes of mechanical stretching, respectively. This trend continued for the next two weeks. Further experimental details are presented in table S4.

### Cuticle and stress response genes are key components of the WT genetic response to mechanical stretching

We used the RNA-seq data for gene counts and FPKM (Fragments Per Kilobase of transcript per Million mapped reads) distributions to compare the global expression pattern between mechanically stretched and un-stretched WT animals. This comparison showed 2484 genes that were significantly altered, with 1351 of those genes being upregulated and 1133 genes downregulated (Fig. 2A). Significance was defined as log2 fold change of more than 1, p value less than 0.05 and at least 1 FPKM level of expression in at least one of the experimental conditions. To further map these early response genes, we performed cluster and gene ontology (GO) analyses (Fig. 2B). Among the most significantly up regulated clusters was animal locomotion, with 212 upregulated genes. The cuticle is composed of specialized extracellular matrix, which protects the animal from external insults by establishing an impervious barrier between the worm and the environment. Indeed 137 genes involved in maintaining cuticle integrity, including 33 genes involved with cuticle development were also up regulated. Stretching *C. elegans* also evoked the upregulation of 164 stress response genes, as well as 68 genes involved in animal defense response. Among the upregulated stress response genes, we also detected 31 genes involved in the response to topologically incorrect proteins (unfolded protein response-UPR) (Fig. 2B, table S1). Downregulated genes included 256 genes involved in reproduction, 38 genes involved in cellular communication and 111 genes involved in gene expression (Fig. 2B and table S1). When comparing the transcriptional response of stretched animals with that of animals that were placed between the two silicone membranes, but not mechanically stretched, we found only a small number of genes in common, 30 genes of the 1351 upregulated when animals are stretched, and 45 genes of the 1133 downregulated mechano-sensitive genes (S2). The expression level of 11 genes was analyzed using qPCR and the results were compared to the transcriptome data (S3). We found a similar direction in the gene expression response to that of the RNAseq data in all tested genes, thus validating the significance of our transcriptome analyses (S3).

**Figure 2:**
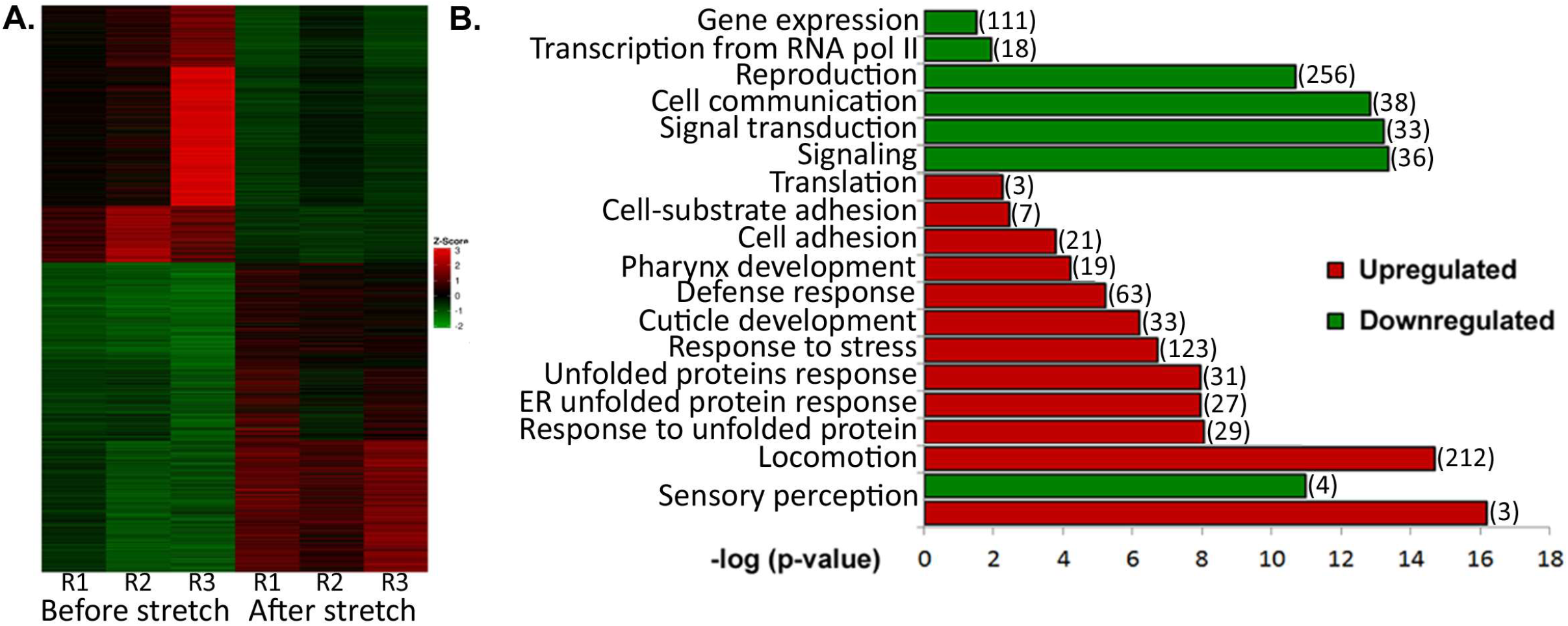
Genetic changes in WT animals following mechanical stretching. (A) Heat map representing genes that were significantly (p-value < 0.05) upregulated (red, n=1364) or downregulated (green, n=1130) in WT animals following the application of the mechanical strain. R = repetition of the experiment (B) (–log) of the p-value score of gene clusters involved in biological processes up regulated (red) and down regulated (green) following Gene Ontology (GO) Enrichment Analysis (http://geneontology.org/page/go-enrichment-analysis). (#) represents the number of genes in each cluster.

Of particular interest is the up regulation of a large number of the hedgehog signaling pathway genes (n=30, Table S1). This discovery prompted us to test the importance of an intact hedgehog pathway to the ability of animals to recover following a mechanical insult. We tested one of the two known hedgehog receptor orthologs in *C. elegans*, *ptc-3,* which in the transcriptome data was shown to be 2 folds upregulated following mechanical stress application. WT and *ptc-3* (RNAi) animals were placed between two silicone membranes, stretched for 5 min to 140% of their original surface area and scored for their ability to survive this stretching (Fig. 3D). Downregulation of *ptc-3* led to a steep and significant drop in animal survival already during the same day of stretching, as compared to stretched animals treated with an empty vector (EV). Un-stretched animals treated with *ptc-3* RNAi or EV did not show mortality during the 1st day (Fig. 3D). We concluded that *ptc-3*, and potentially intact hedgehog signaling, are required for *C. elegans* to survive mechanical stress. We also looked at the effect of several genes known to be involved in mechano-sensation on the ability of *C. elegans* to recover from mechanical stretching, including *pezo-1*, an ortholog of the human piezo type mechanosensitive ion channel, which was found in our analysis to be 2.37 folds upregulated in response to mechanical stretching (Fig. 3C), *mec-15* (Fig. 3B, 1.87 folds downregulated), an F-box protein responsible for proper function of all six *C. elegans* touch neurons and *mec-3* (Fig. 3A, un-affected), a transcriptional regulator expressed in touch neurons on animal survival post stretching. Analysis of animal survival a few hours following mechanical stretching did not show any significant changes between animals treated with EV and animals downregulated for *mec-3*, *mec-15* or *pezo-1* (Fig.3A-C). This implies that these genes are not involved in the short-term survival of animals following mechanical insults. One day following stretching, *mec-15* and *pezo-1* (RNAi) animals have only slightly reduced survival (∼88%) as compared to *mec-3* (RNAi) and EV animals (∼94% and ∼93%, respectively) (Fig.3A-C), indicating they may be involved in the long-term survival of animals following mechanical stress.

**Figure 3:**
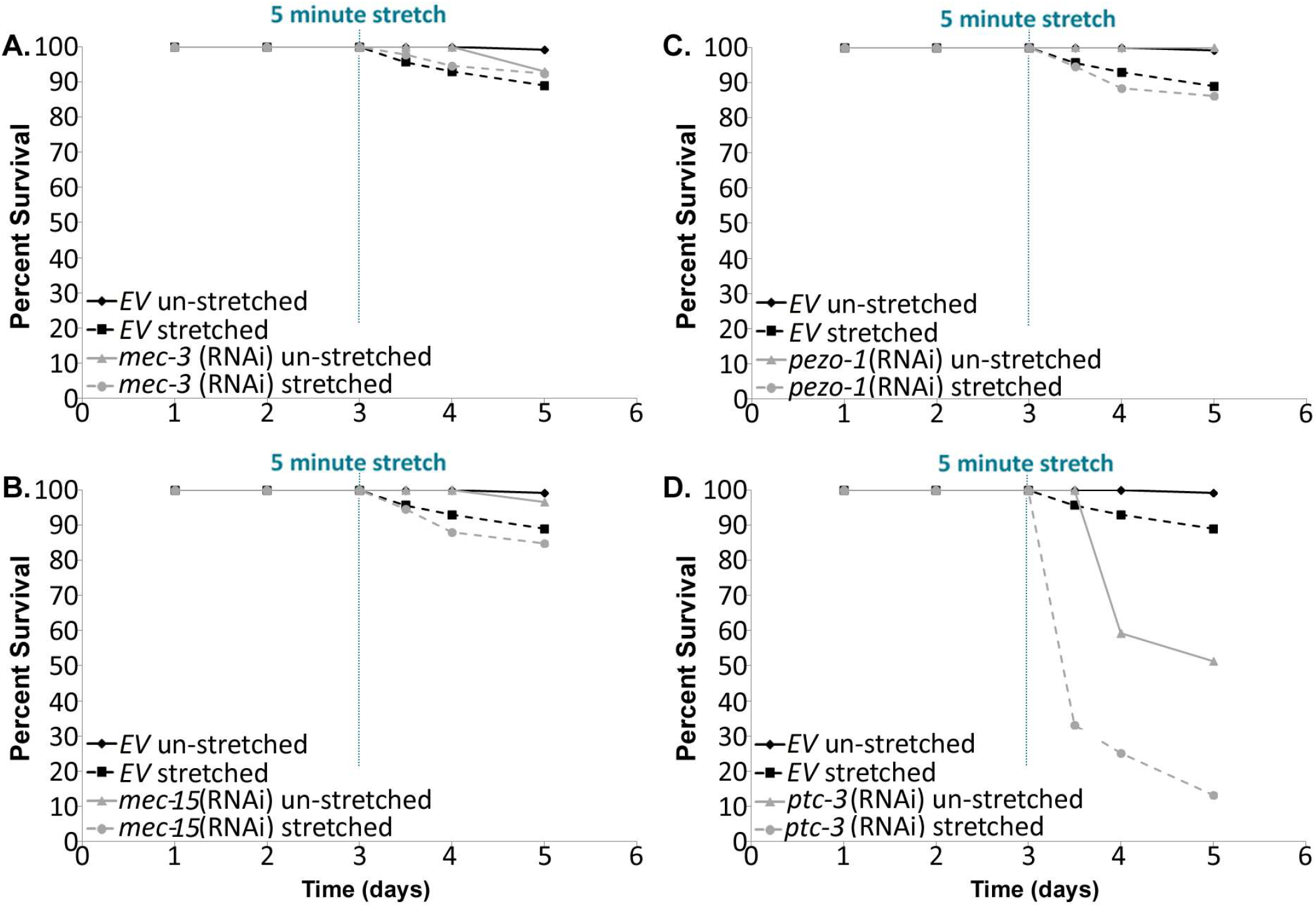
Survival curves for animals at day 2 of development (L4), following treatment with either (A) *mec-3,* (B) *mec-15,* (C) *pezo-1* or (D) *ptc-3* (RNAi) before and after 40% stretching for 5 min. WT worms were fed with empty RNAi vector (EV) and served as controls. Within few hours of stretching, roughly 5% lethality was detected in WT animals treated with EV (dashed black, n=411). *mec-3*, *mec-15* and *pezo-1* (RNAi) treatment did not affect lethality significantly at this time point (dashed grey, n= 91, 92 and 385, respectively). *ptc-3* (RNAi) treatment however induced a statistically significant sharp increase in animal lethality (dashed grey, n=376), elevating it to nearly 70%. No significant lethality was observed in the un-stretched worms treated with *EV* (black, n=120). Further experimental details are presented in table S4.

To test whether the genetic response observed following stretching is unique to mechanical stress, we compared it with other previously reported types of stressors; heat shock, hypoxia and hypercapnia (S4). To our surprise, the majority of genes upregulated (1058 out of 1351), as well as downregulated (980 out of 1133) by stretching were not in common with the other types of stress (S4). Among the unique mechanically upregulated genes (n=1058), 77 genes clustered as stress response genes, 65 were cuticle genes, 18 were hedgehog genes and 14 were cytoskeleton genes.

### Cytoskeleton organization and cellular respiration genes are upregulated following strain application in L535P EDMD animals

We next determined how the mechano-sensitive genetic response to strain application is altered in lamin disease conditions. A previous study showed changes of hundreds of gene including key genes and biological pathways that can help explain the muscle specific phenotypes of animals expressing the L535P mutation in the lamin gene [21]. The mechanically-induced transcriptional profiles of WT and L535P animals were compared using a custom-made Python program (S5, also see methods section). We identified 1796 genes that were specifically altered in the mechanically-stretched L535P animals, as compared to mechanically-stretched WT animals. Among those, 758 genes were upregulated and 1038 genes were downregulated. Further GO analyses revealed that the biological processes upregulated in these stretched mutant animals included genes that were clustered to cytoskeleton organization (n=45), muscle organization (n=16), muscle contraction (n=9) and cellular respiration (n=14) (Fig. 4A). We next examined whether the altered genes could represent specific molecular pathways affected by strain application. Using our custom-made interaction fetcher (S6, see also methods section), we detected genes whose interaction profile and mechano-induced expression levels matched genetic pathways involved in mitochondrial unfolded protein stress response (Fig. 4Ba) and animal motility (Fig. 4Bb-f).

**Figure 4:**
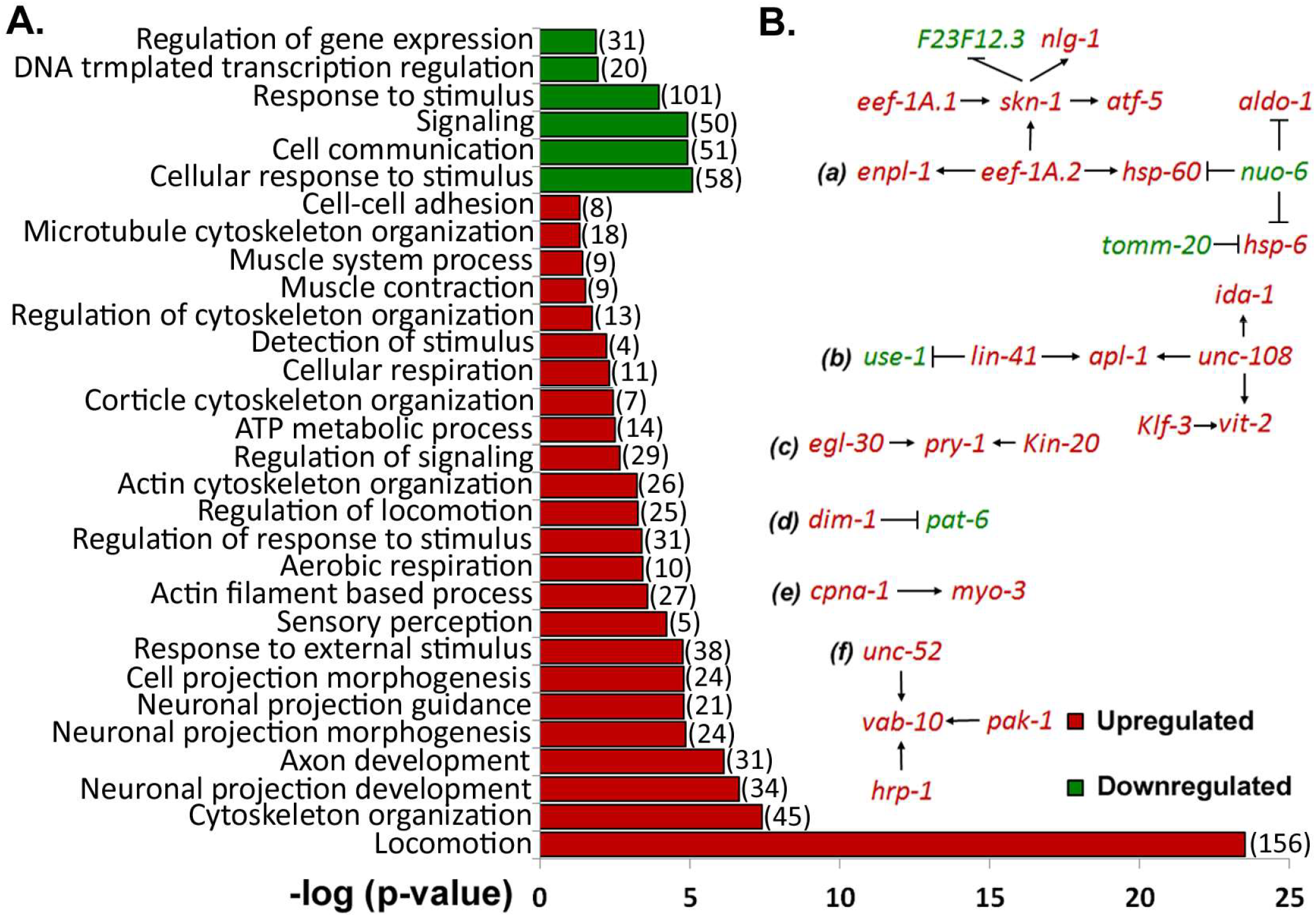
Uniquely altered genes in L535P animals following mechanical strain application. (A) (–log) of the p-value score of upregulated (red) and downregulated (green) gene clusters following Gene Ontology (GO) Enrichment Analysis. Displayed are biological processes affected by the application of mechanical strain only in L535P animals. (#) represents the number of genes in each cluster. (B) Molecular pathways uniquely activated in mutant L535P animals following strain application. These pathways involve mitochondrial unfolded protein stress response (a) and animal motility (b-f). Genes indicated in red are upregulated, while genes indicated in green are downregulated.

The changes caused by the expression of the L535P mutation can be explained by a compensatory mechanism in which genes that were negatively affected merely by the expression of the L535P mutation [21], are now upregulated when these animals are being stretched. Indeed, comparing genes downregulated by the L535P lamin mutation in un-stretched animals, with genes uniquely upregulated in the mechanically stressed L535P expressing animals, revealed 387 genes in common, accounting for 51% of the uniquely mechano-upregulated genes in the L535P expressing animals (Table S2). This compensatory effect included genes involved in establishing proper muscle activity and animal motility, which are impaired in un-stretched L535P EDMD animals (Fig. 5) [21,22]. For example, actin isoforms 1-4, which are similar in structure, and *tni-3*, encoding for one of the four *C. elegans* troponin I genes, are required to coordinated locomotion, as well as, proper muscle structure and contractions. These genes were downregulated in un-stretched L535P animals but upregulated as a result of stretching. *pfn-1* (profilin), which binds actin and affects the structure of the cytoskeleton; *cmd-1* encoding the *C. elegans* calmodulin and *dyn-1* (ortholog of the dynamin GTPase), which is involved in endocytosis and synaptic vesicle recycling and required for proper animal locomotion, were also similarly affected. In our previous analysis, we have shown that genes involved in cellular respiration and ATP metabolism were negatively affected by the expression of the L535P lamin mutation and mitochondria of these animals were mis localized [21]. Following stretching, we could detect several of these mitochondrial genes upregulated; *ucr-1* which is an ortholog of human PMPCB (peptidase, mitochondrial processing beta subunit) and UQCRC1 (ubiquinol-cytochrome c reductase core protein I); *asb-2* and *F58F12.1* encode subunits of complex V of the mitochondrial respiratory chain; *sdha-1, sdha-2* and *mev-1*, all encoding subunits of mitochondrial complex II; *H28O16.1*, an ortholog of the mitochondrial ATP synthase alpha subunit and *W09C5.8* which encodes an ortholog of the cytochrome c oxidase (COX) subunit IV (COX IV) (Fig. 5, Table S2).

**Figure 5:**
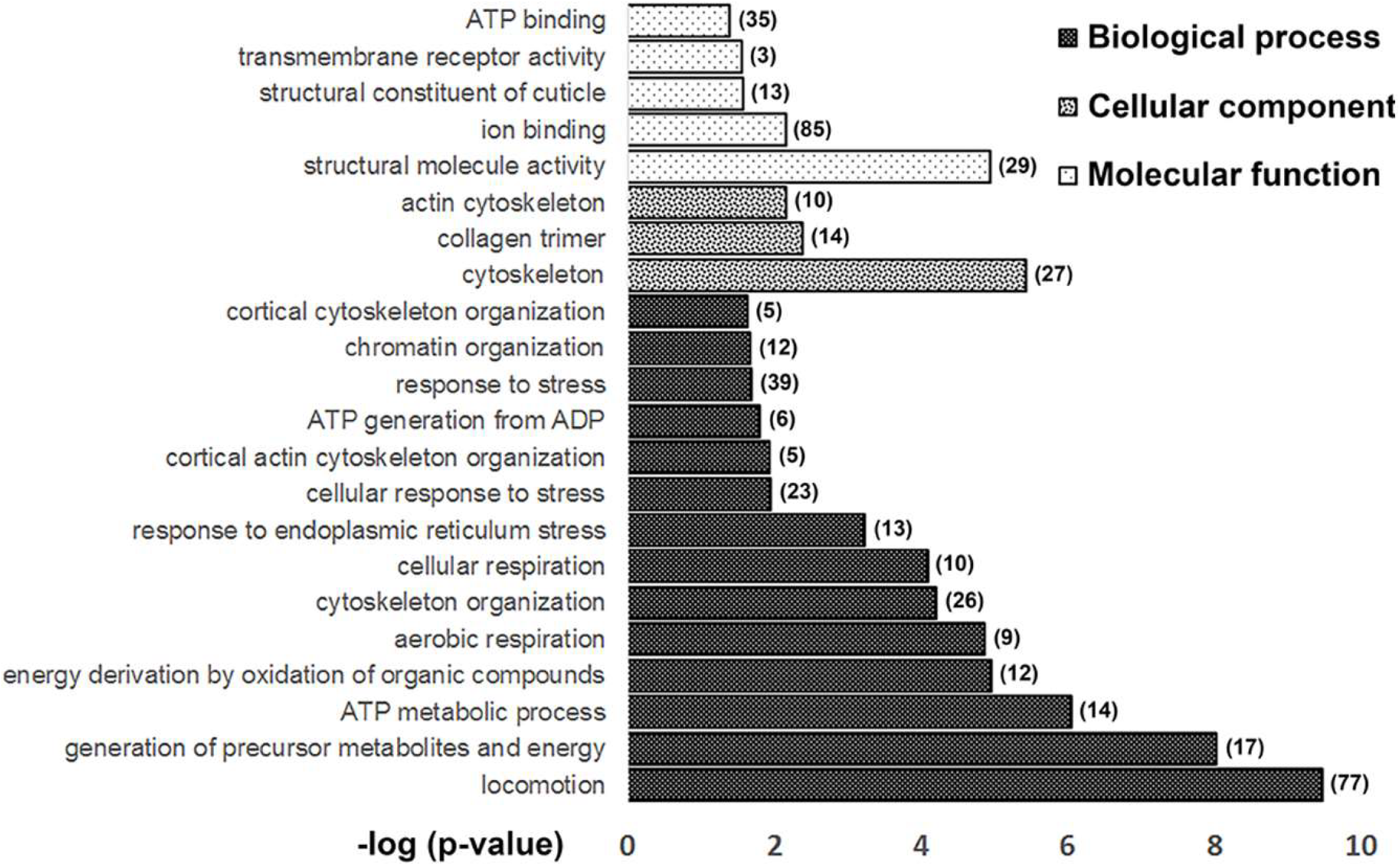
Compensatory genetic response to the application of mechanical strain in L535P animals. (–log) of the p-value score of gene clusters following Gene Ontology (GO) Enrichment Analysis. Shown are clusters of genes that are both uniquely upregulated in stretched L535P animals and down regulated in un-stretched animals expressing the L535P lamin mutation.

Alongside the changes in mechanically-induced gene expression profiles caused by the L535P EDMD-linked mutation, we also identified 1203 genes that were altered in a similar way in both WT and L535P animals; 594 upregulated genes and 609 downregulated genes. GO analyses revealed 69 genes involved in the biological processes of stress response, 25 genes involved in ER un-folded protein response and 81 genes involved in maintaining cuticle integrity, including 22 genes involved with cuticle development that were upregulated in both stretched WT and L535P animals (Fig. 6A). A large number of the hedgehog genes were also un-affected by the L535P mutation (n=17, Table S3).

**Figure 6:**
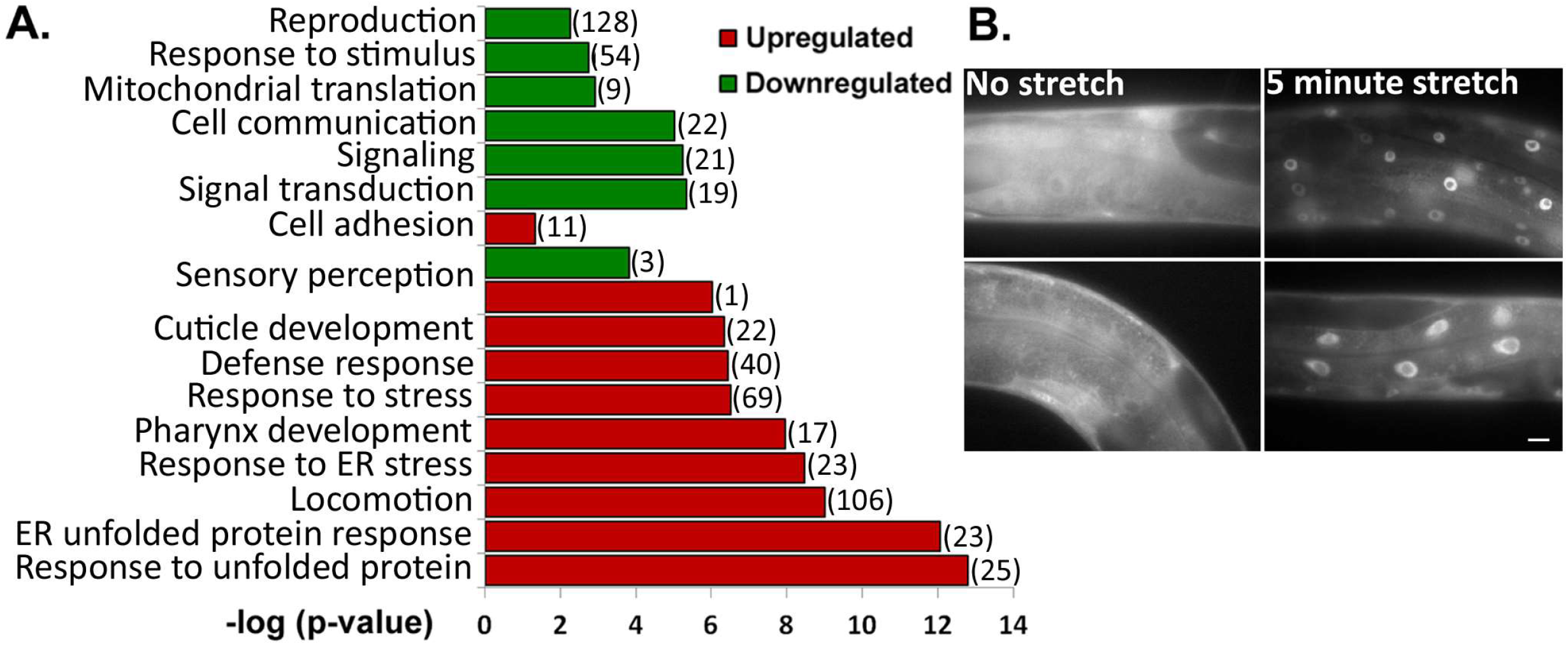
Common changes in mechanically induced transcription between WT and L535P animals. (A) (–log) of the p-value score of upregulated (red) and downregulated (green) genes clusters to biological processes following Gene Ontology (GO) Enrichment Analysis. Shown are genes similarly altered by mechanical strain application in both WT and L535P animals. (#) represents the number of genes in each cluster. (B) *daf-16*::GFP localizes to the nucleus following 5 minutes of mechanical stretching. Scale bar-10μm.

## Discussion

We have analyzed the global transcriptional response to the application of mechanical stress in whole living *C. elegans* animals. For this, we established an experimental system, which allows simultaneous analysis of hundreds of living worms (S1). The use of this experimental setup enabled the identification of a large number of mechano-sensitive genes, spanning multiple tissue types and genetic pathways. This in turn, gave a multi-dimensional picture of the genetic response of animals to mechanical stress. To date, such experiments have been done only on isolated cells or muscle fibers [23–25], limiting the detection of relevant mechano-induced genetic response to that occurring in specific cell types or tissues isolated from their environment.

Previous studies of the response of *C. elegans* to stress have shown that different types of stressors cause *daf-16* to be relocated to the nucleus, where it induces the transcription of stress response genes, promotes dauer formation and reduces fecundity [26]. Indeed, when animals are subjected to mechanical strain, we detect a rapid shift of *daf-16* from the cytoplasm to the nucleus (Fig. 6B). We did not detect, however, a corresponding increase in *daf-16* transcription. These observations may indicate that the initial response of animals to mechanical stress is initiated by the movement of some of the readily available pool of cellular proteins, such as *daf-16*, and their subsequent activity at their target locations. Following our RNA-seq analysis, the mechanically stressed WT animals showed a distinct up regulation of genes involved in animal stress response, as well as downregulation of reproduction genes. Comparing our mechanically induced genetic response to that of other *C. elegans* stressors showed that most mechanically induced genes were different from genes altered in response to heat shock, hypoxia and hypercapnia (S4). Among the 164 identified stress response genes found to be upregulated in our analysis, 77 were unique to mechanical stress response. In addition, genes involved in cuticle development and integrity were specifically upregulated in the mechanically stressed animals as compared to other types of stressors (Fig. 2, Table S1). To verify that the observed genetic response was caused by the application of mechanical stretching, and not by the mere placement of animals between two silicone membranes, we compared the transcriptional changes occurring when animals are placed between the two membranes, with and without mechanical stretching. This analysis showed a small number of genes in common (∼2% of the upregulated genes and ∼4% of the downregulated genes, S2) indicating that the mechano-sensitive transcriptional changes indeed resulted from the application of mechanical stretching. These findings indicate that a first line of defense in animals subjected to mechanical stressors is specific and is mainly comprised of their ability to alter transcription and signaling to upregulate genes that boost the resilience of the cuticle and the cytoplasmic filament networks, presumably to maintain and provide additional support to this mechanically resilient protective structure. In addition, the relatively rapid transcription upregulation of these genes indicates that *C. elegans* are genetically preprogrammed to be able to defend against mechanical insults. Our analysis also revealed transcriptional changes in genes involved in molecular pathways such as WNT, TGFβ and AMPK (Table S3). These signaling pathways were previously shown to change in response to external stress application on cultured cells (3-5), providing additional confirmation to our results.

Of particular interest is the observation that mechanical stress application increases the transcription of a large number of genes involved in the hedgehog (Hh) signaling pathway. We further demonstrate that this pathway is required for animal recovery from mechanical stress by showing that inhibition of a key gene in this pathway, the PATCHED gene homologue *ptc-3*, at the L4/young adult stage prevented the ability of the animals to recover from stretching. PTC-3 is required for normal molting from L1 to L3, as well as for normal growth. It is therefore possible that potential defects in the animal cuticle contribute to the inability of *ptc-3* (RNAi) animals to recover from mechanical stress. Indeed, 30 of the 65 cuticle genes that were found to be mechanically upregulated in our analysis are required for proper molting. When worms are stretched, muscles are exposed to a high degree of strain, which could induce muscle tearing and injury. Such defects in muscles may also contribute to the inability of *ptc-3* (RNAi) animals to recover from mechanical stress. This hypothesis is supported by previous studies performed on mice, which demonstrate that the hedgehog signaling pathway is activated upon skeletal muscle injury [27,28]. Ptc-1 in particular is expressed in the adult rodent heart, upon ischemic injury, the hedgehog pathway is activated, and sHh and Ptc-1 expression is upregulated [29,30]. These findings are in line with the overall stretch-induced upregulation of hedgehog genes observed in our analysis (Table S1).

It has been shown that muscle nuclei of *C. elegans,* expressing the L535P EDMD-linked mutation, employ an aberrant mechanical response, which, when corrected, rescued the disease phenotypes [20]. Moreover, this mutation causes global changes in transcription of genes involved in body wall and pharyngeal muscle function and structure [21]. Interestingly, in L535P animals subjected to mechanical stretching, many genes involved in proper muscle function, cytoskeletal organization and cellular respiration were uniquely upregulated (Fig. 4). In the disease situation, the muscle cells experience an increased degree of stress, due to their impaired structure and function. It is conceivable that to better handle the destructive effects of the added stress; genes that help to improve muscle function and oxygenation are activated. Indeed, a large number of the genes that were uniquely upregulated in mutant animals following stretching, were shown to be negatively affected by the expression of the L535P mutation in a previous study (S6, table S2) [21].

Along-side the changes in animal’s response to mechanical stimulation caused by the L535P mutation, several key pathways remained un-affected (Fig. 6). The activation of genes involved in cuticle development indicates the importance of this structure in maintaining animal integrity and providing protection from external insults in both WT and disease situations. It is important to note that cuticle genes were also up regulated in animals expressing the L535P mutation, which did not undergo mechanical stretching [21]. The fact that the mechano-sensitive transcriptional response of the majority of hedgehog genes was not altered by the L535P mutation, including *ptc-3*, further highlights the importance of this molecular pathway in mediating animal response to mechanical stress. Recent studies performed on patient with heart failure, demonstrated that decreasing the mechanical load experienced by myocardial tissue, which is achieved by the implantation of a left ventricle assist device (LVAD), decreased markers of ER UPR [31]. These findings are consistent with UPR being one of the key pathways activated in both WT and mutant animals following the application of mechanical strain.

There are many lamin based muscle diseases with varying degrees of severity and phenotypes. These diseases are caused by a large number of globally expressed lamin mutations. Understanding the WT genetic response to mechanical stress, as well as how the transcriptional mechano-sensitive profile changes as a result of the expression of different EDMD mutations can serve as a future direction towards finding the common pathways affected in these diseases both in rest and under mechanical strain. This insight can then be implemented to develop treatment options for these types of diseases.

## Acknowledgments

We thank Liran Carmel and Alon Zaslaver for a critical reading of the manuscript. This work was funded by grants from the Israel Science Foundation (ISF, 785/15), the German Israel Foundation (GIF I-1289-412.13), the Niedersachsen-Israeli Research Cooperation and by the European Cooperation in Science and Technology (COST CA grant 15214, EuroCellNet).

## Author Contributions

N.Z.S. and Y.D. performed experiments, N.Z.S. and Y.G. conceived experiments, Y.D. programming, N.Z.S. and Y.D. wrote the manuscript, Y.G. secured funding.

## Materials and Methods

### Strains

Strain maintenance was performed under standard conditions as described previously [32]. The Bristol strain N2 was used as a standard WT while the YG499 ([*baf-1*p::GFP::*lmn-1* L535P *unc119*(+)]; *unc-119*(ed3)) strain was used as an EDMD disease model. TJ356 ([*daf-16*p::DAF-16::GFP + *rol-6*(su1006)) was used for tracking DAF-16 localization. Strains were out-crossed at least 3 times to ensure a clean background.

### *C. elegans* population stretching

To map early changes in global mechano-sensitive transcription in live animals, RNA had to be extracted from a large number of *C. elegans* animals that are subjected to simultaneous mechanical strain. N2 and YG499 worms were bleached in 0.06 M NaOH, 0.5% sodium hypochlorite solution, and the resulting embryos were synchronized by shaking in M9 buffer (3g KH2PO4, 6g Na2HPO4, 5g NaCl, 1ml 1M MgSO4, H2O to 1 liter. Sterilize by autoclaving) over night at (20°C) and subsequently moved to NGM plates. Late L4 animals were collected and washed 3 times with M9. After the last wash, most of the M9 was removed and roughly 300 worms were suspended in 0.1% poly-lysin (v/v), to facilitate worm adhesion (S1). Tiny drops of the worm containing solution were placed on a silicone membrane (Silicone sheeting 005″ NRV G/G 40D 12″×12″, Specialty Manufacturing, MI 48603-3440). This membrane was subsequently covered with a second silicone membrane making sure not to trap any air in between. The two silicon membranes were mounted on a strain application device [20] and stretched in a multi-directional manner to about 140% of their original surface area for a time period of 5 min (red arrows, S1). Once the strain has been eliminated, the two membranes were separated, and worms were washed from the membranes using M9. Following a 40 minute recovery period, *C. elegans* were fully recovered with respect to their viability and motility (Fig. 1). Un-stretched N2 and YG449 worms were suspended in 0.1% poly-lysin (v/v) for the duration of the stretching procedure; than washed with M9, to account for genetic changes that may occur as a result of the suspension in 0.1% poly-lysin. Each experiment was repeated three times.

### RNA isolation

Performed as previously described in [21]. In short, 40 min after the release from stretching, stretched and un-stretched N2 and YG449 animals were treated with BIO TRI RNA (Bio-Lab Itd., Cat. #90102331) and kept overnight at (−80°C), subjected to 3 rounds of crushing and freezing in liquid nitrogen, followed by a 12,000xg centrifugation for 15 minutes at 4°C. Supernatant was transferred to a new tube and treated with chloroform. The aqueous transparent phase was transferred to a new tube, mixed with isopropanol and incubated overnight at −20°C. 75% ethanol was added to the pellet following a 12,000 x g centrifugation at 4°C for 30 min. Pellets were air-dried on ice and eluted using PCR grade DDW. RNA concentration was measured using a Bio-Analyzer machine (Agilent Technologies, Inc., serial #DE54700884, firmware C.01.069, type G2939A) and frozen at −80°C until use.

### Complementary DNA (cDNA) preparation

DNase treatment (RQ1 RNase-Free DNase, Promega Corporation, cat. #M6101) and cDNA synthesis (M-MLV RT kit, Promega Corporation, cat. #M170A) were performed according to manufacturer’s specifications [21]. Samples were stored at −80°C until use.

### RNA sequencing

Sequencing of the cDNA libraries was performed using the NextSeq 500 High Output kit (75 cycles) #FC-404-2005 Illumina by the Center of Genomic Technology at the Hebrew University.

### Bioinformatics analysis

Sequence data processing was performed essentially as described in [21]. In short, Quality control was performed on the raw reads with FastQC (v0.11.2, http://www.bioinformatics.babraham.ac.uk/projects/fastqc/). Reads were filtered and quality-trimmed at both ends, using in-house Perl scripts. SL1 and SL2 sequences were removed from the 5’ end using cutadapt (version 1.7.1, http://cutadapt.readthedocs.org/en/stable/). The adapter sequence was then similarly trimmed from the 3’ end. The remaining reads were filtered for reads that were shorter than 15 nucleotides as well as reads with very low quality, using the fastq_quality_filter program of the FASTX package (version 0.0.14, http://hannonlab.cshl.edu/fastx_toolkit/).

The processed fastq files were aligned to the *C. elegans* genome (version WBcel235) using TopHat (v2.0.13). The Cufflinks package (v2.2.1) was used for quantification, normalization and differential expression. The cummeRbund package (version 2.8.2) and in-house R scripts were used to visualize results. Comparison of global expression, as well as background expression level estimation between samples was measured using FPKM distributions. Differential expression was calculated with cuffdiff, using a count threshold > 15 for statistical significance testing. Statistically significant differential expression was defined as genes with at least 1 FPKM level of expression in at least one of the experimental conditions and a Benjamini-Hochberg-adjusted P value (q-value) less than 0.05. Results were visualized in R with in-house scripts based on the cummeRbund code.

### Cluster analysis

Significant genes (−1>log2 fold change>1, p-value<0.05) were clustered using DAVID Bioinformatics Resources 6.7 (https://david.ncifcrf.gov/tools.jsp) and GO Enrichment Analysis | Gene Ontology Consortium annotation tool (http://geneontology.org/page/go-enrichment-analysis).

### Comparative and interaction analysis

Gene groups of interest were compared to one another using a custom-made Python program which can be downloaded from (https://bitbucket.org/Jehudith/listcomparison/src) (S5). To compare the mechanically induced stress response to other stress responses, previously reported databases were used; Heat shock data set 1 was retrieved from https://www.ebi.ac.uk/arrayexpress/search.html?query=%2C+E-MTAB-5753, Heat shock data set 2 was retrieved from [33], Hypoxia data set [34] and hypercapnia data set from [35]. Interaction analysis was performed using a custom-made Python program which displays all interactions of a certain gene, as well as the specific gene of interest’s interacting partners within a certain cluster. The program can be downloaded from (https://bitbucket.org/Jehudith/interactionsearcher) (S6).

### Quantitative PCR analysis

Quantitative amplification was performed according to manufacturer’s specifications using KAPA SYBR® FAST qPCR Kit Master Mix (2X) Universal (Sigma, cat. #07959397001). 12 separate RNA isolations were analyzed, 6 from WT (un-stretched and stretched in triplicates) and 6 from YG449 animals (un-stretched and stretched in triplicates). Rotor-Gene™ machine (Corbett Life Science, Sydney, Australia, Model RG-6000 #R 120726) with software version 1.7 (Build 87) was used for analysis. Primers were designed according to qPCR standards [36] using ApE (v2.0.3.7, Copyright © 2003-2009 by M. Wayne Davis). The *C. elegans* housekeeping gene *pmp-3* was used as control. (Primer sequences are reported in S7)

### Lifespan Assay

N2 animals were bleached and placed on feeding plates seeded with empty vector (EV), *mec-3, mec-15, pezo-1* or *ptc-3* (RNAi). Late L4 animals were washed from the plates and stretched for 5 min as described (See *C. elegans* population stretching). Animals were washed from the membranes and placed back on the respective feeding plates seeded with either EV, *mec-3, mec-15, pezo-1* or *ptc-3* RNAi bacteria, respectively. Scoring was initially performed a few hours after the application of mechanical stress, as well as during the week following stretching. For survival following different stretching times, N2 animals were bleached, synchronized and placed on OP50 plates. Late L4 animals were washed from the plates and stretched for 5, 15 and 30 minutes as described (See *C. elegans* population stretching). Animals were washed from the membranes and placed back on OP50 plates. Scoring was initially performed roughly an hour after the application of mechanical stress, as well as every day during the following two weeks. Live animals were scored as responsive to gentle prodding with a platinum wire. Animals were moved to new plates every other day while laying eggs. Survival plots were generated using OASIS [37]. Summary of experimental data is depicted in Table S4.

### Fluorescence microscopy

Late L4 C. *elegans* expressing *daf-16* fused to GFP were stretched between two silicone membranes for 5 minutes. Worms were washed from the membranes and placed on 2% agar slides. *Daf-16* localization was scored in stretched versus un-stretched animals. Images were acquired using an ORCA-R2 camera (Hamamatsu Photonics) mounted on a Zeiss Axioplan II microscope equipped with epifluorescence.

## Supporting information

**Figure S1:**
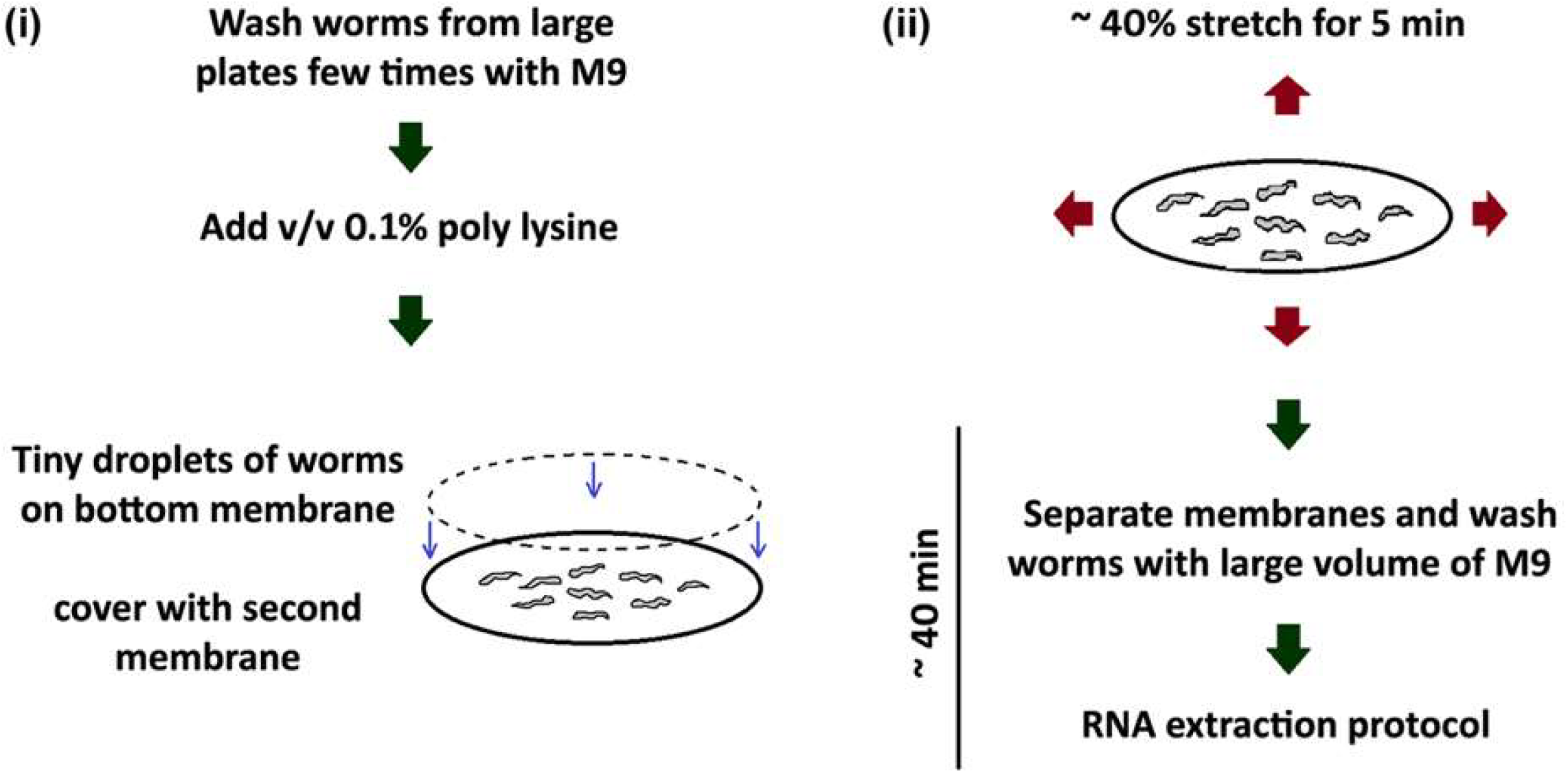
The experimental system. To simultaneously analyze hundreds of worms, tiny drops of synchronized late larval stage (L4) animals in 0.1% poly-lysin solution (v/v) were placed between two silicone membranes. The membranes were then stretched for 5 minutes, resulting in a 40% increase of the initial surface area of the worms. After relaxation, the membranes were separated and worms were allowed to recover for 40 min, and collected for RNA isolation. The RNA was then used to prepare cDNA libraries. Each experiment was repeated 3 times.

**Figure S2:**
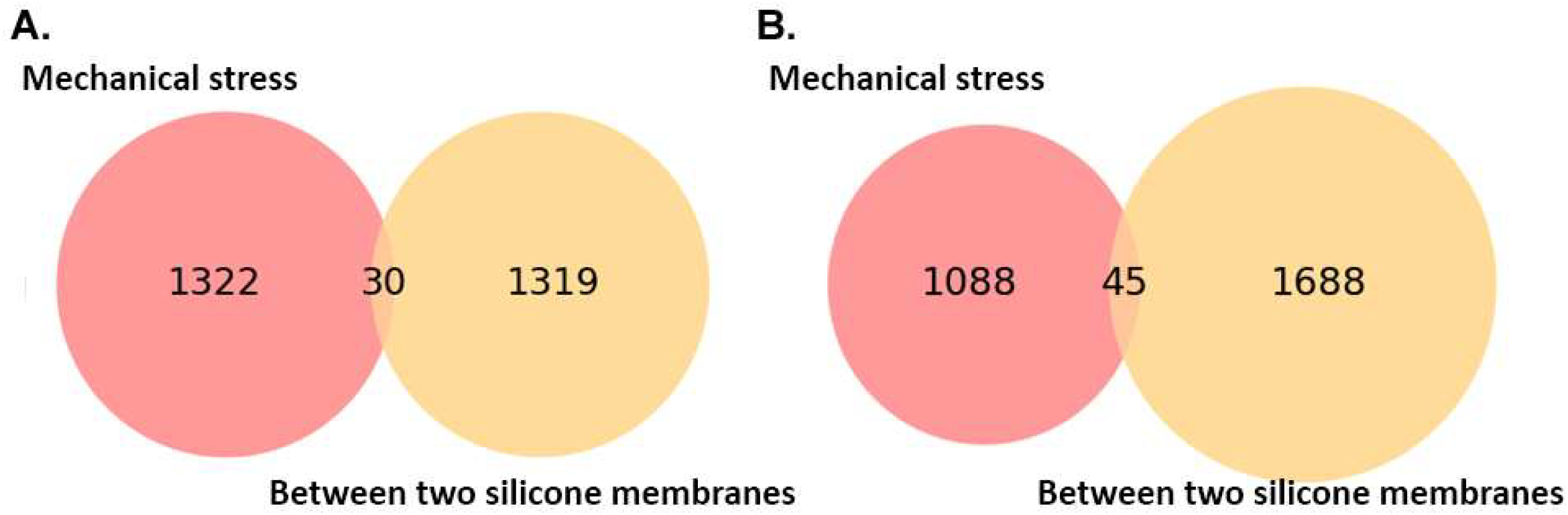
The mechanically induced genetic response is caused by applying mechanical stretching. The list of genes altered by placing worms between two silicone membranes was compared with our list of stretching dependent mechanically induced genes and showed a small overlap of genes between the two, identifying 30 common upregulated genes (A) and 45 common downregulated genes (B).

**Figure S3:**
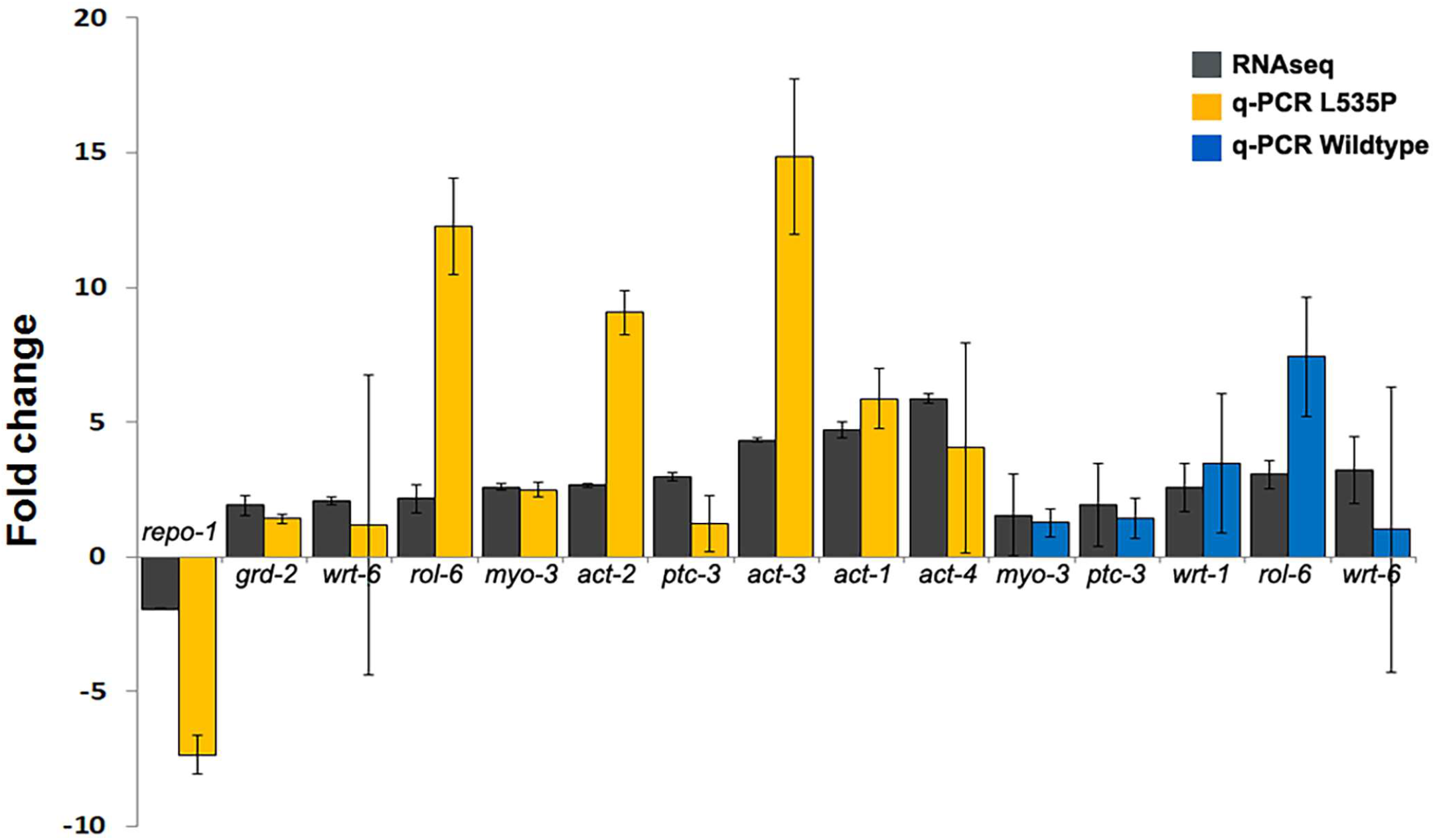
Quantitative PCR verification. Fold change comparison between RNA sequencing (RNAseq) experiments and quantitative PCR (qPCR) experiments. Error bars represent SEM. *P values* of the fold change between qPCR and RNAseq for L535P animals (yellow); *repo-1* (0.04); *grd-2* (0.11); *wrt-6* (0.09); *rol-6* (0.06); *myo-3* (0.0005); *act-2* (0.003); *ptc-3* (0.24); *act-3* (0.012); *act-1* (0.046); *act-4* (0.06) and wild-type animals (blue); *myo-3* (0. 3); *ptc-3* (0.7); *wrt-1* (0.02); *rol-6* (0.58); *wrt-6* (0.42).

**Figure S4:**
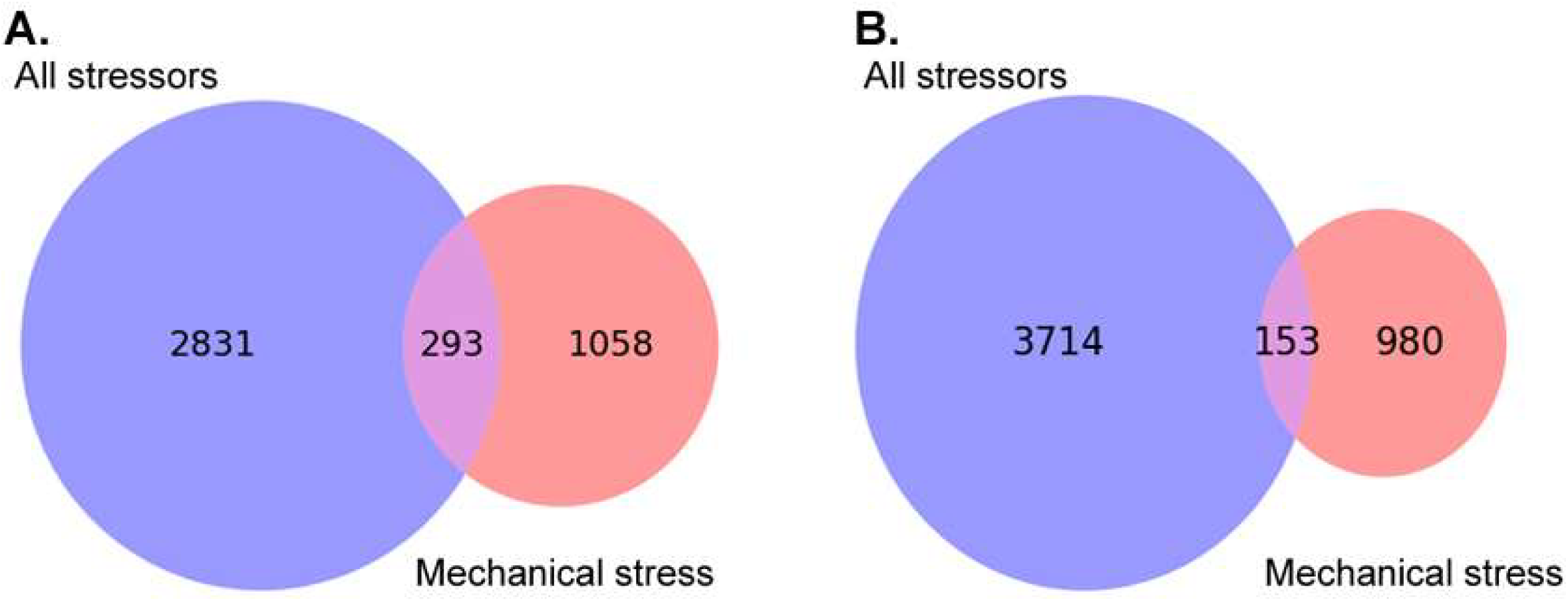
The majority of the mechanically induced genetic response is unique compared to other stressors. Existing databases of known stressors (All stressors) which includ heat shock (data set 1: https://www.ebi.ac.uk/arrayexpress/search.html?query=%2C+E-MTAB-5753, data set 2: [33] Hypoxia [34] and hypercapnia [35] were unified and duplicated genes were removed. The list of unified genes was compared with our list of mechanically induced genes, identifying 1058 unique upregulated genes (A) and 980 unique downregulated genes (B) involved in the response to mechanical stress.

**Figure S5:**
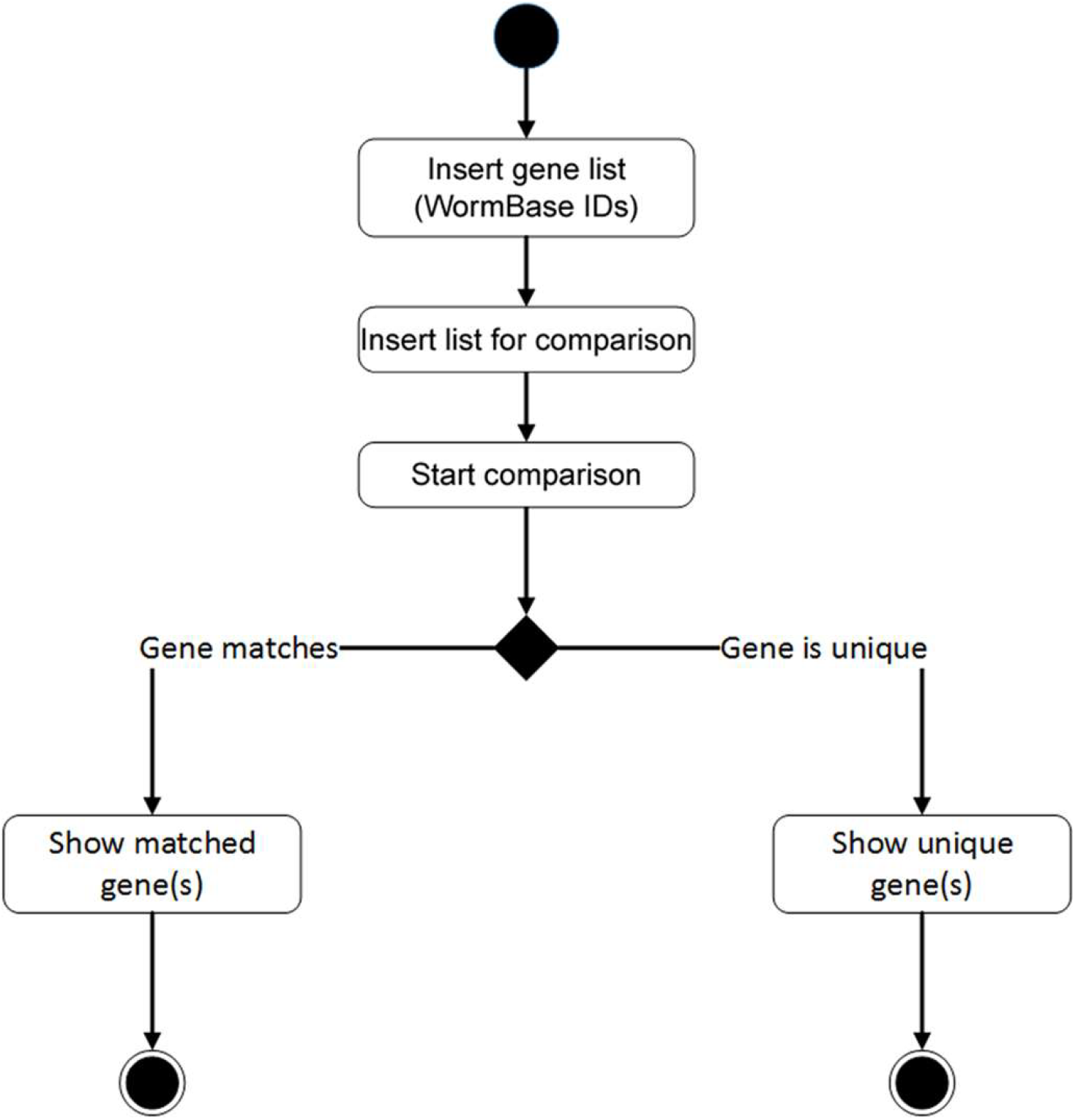
Principles of the custom comparative Python program. To find either common or unique genes of two different lists a custom-made Python program (version 3.3.5) was used. This program enables the user to insert an initial list of WormBase IDs (each ID in a separate line), followed by the list to which the comparison will be made. The program filters all duplicate WormBase IDs and then compares the two lists; the output can be chosen as either a list of matched genes or of unique genes. This program can be viewed and downloaded from https://bitbucket.org/Jehudith/listcomparison/src

**Figure S6:**
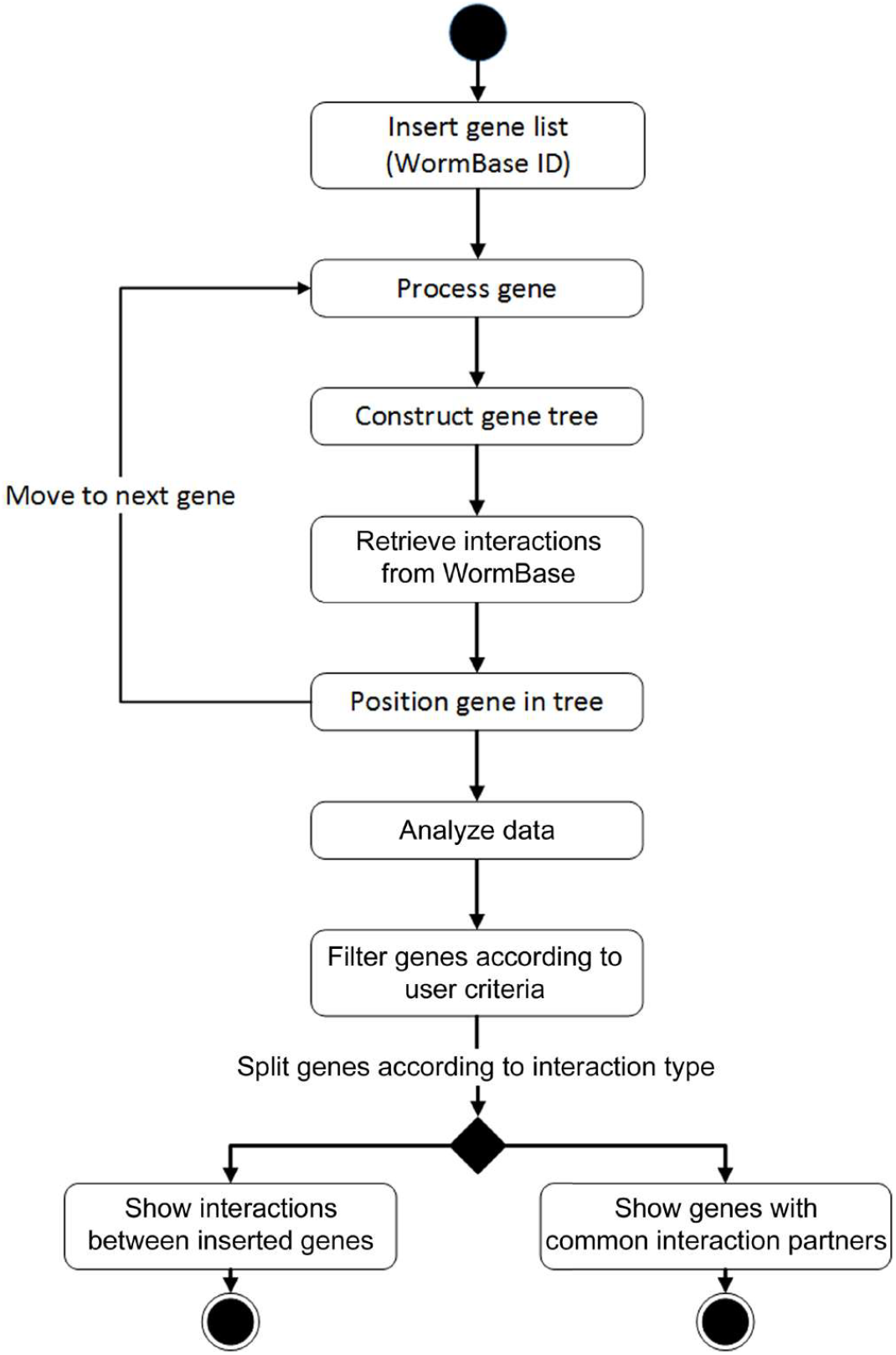
Principles of the custom interaction fetcher Python program. A Python program InteractionSearcher was constructed to find the interacting partners of a given gene, as well as interacting partners within a certain gene list or gene cluster. The program receives WormBase IDs (each ID in a separate line) and retrieves gene interaction information from WormBase. In order to filter the desired interaction type, the program asks the user to filter a specific interaction. The user receives two inputs; interactions within the inserted gene list and genes with common interacting partners. This program can be viewed and downloaded from https://bitbucket.org/Jehudith/interactionsearcher.

**Figure S7:**
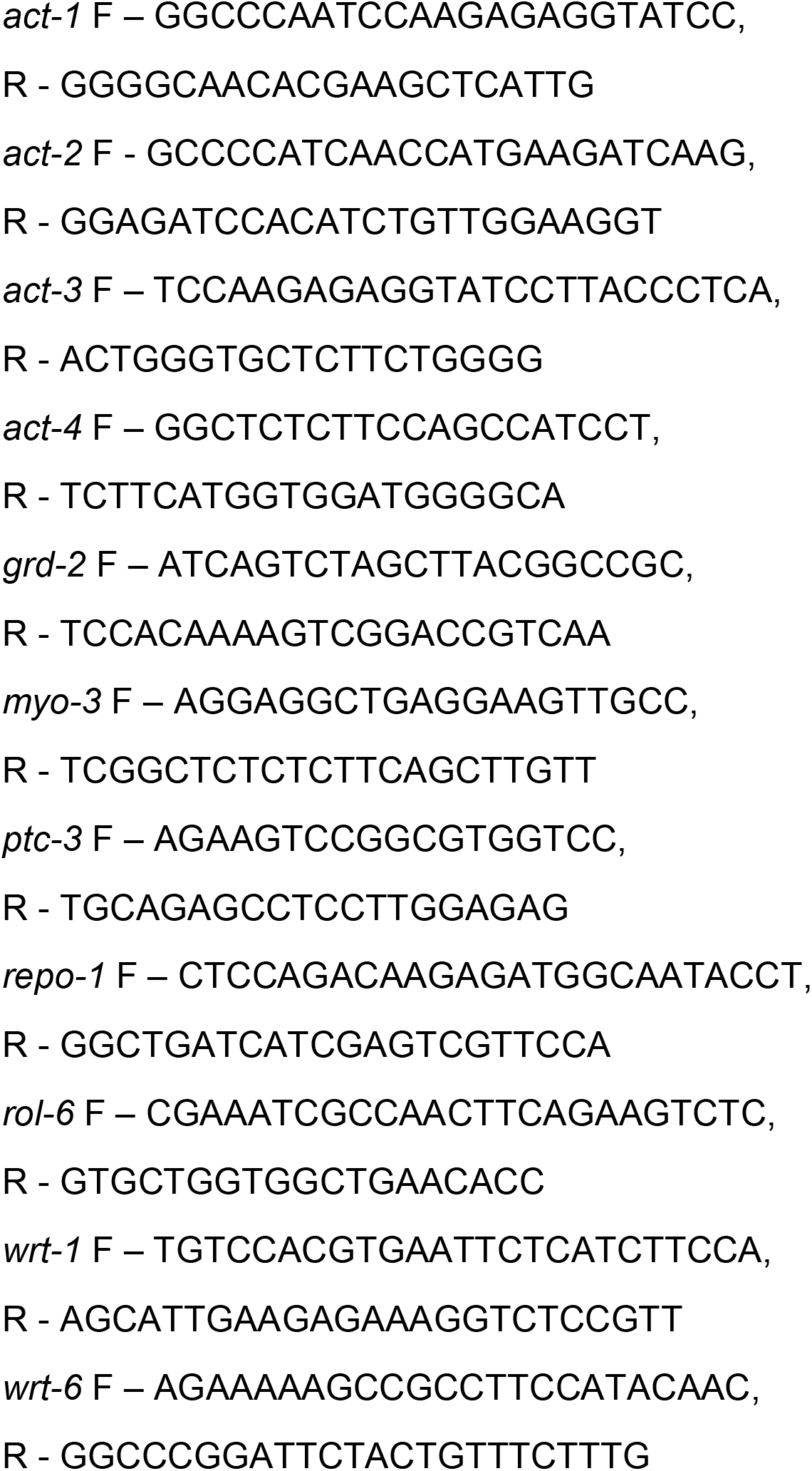
qPCR primer sequences used in this study.

**Table S1:**
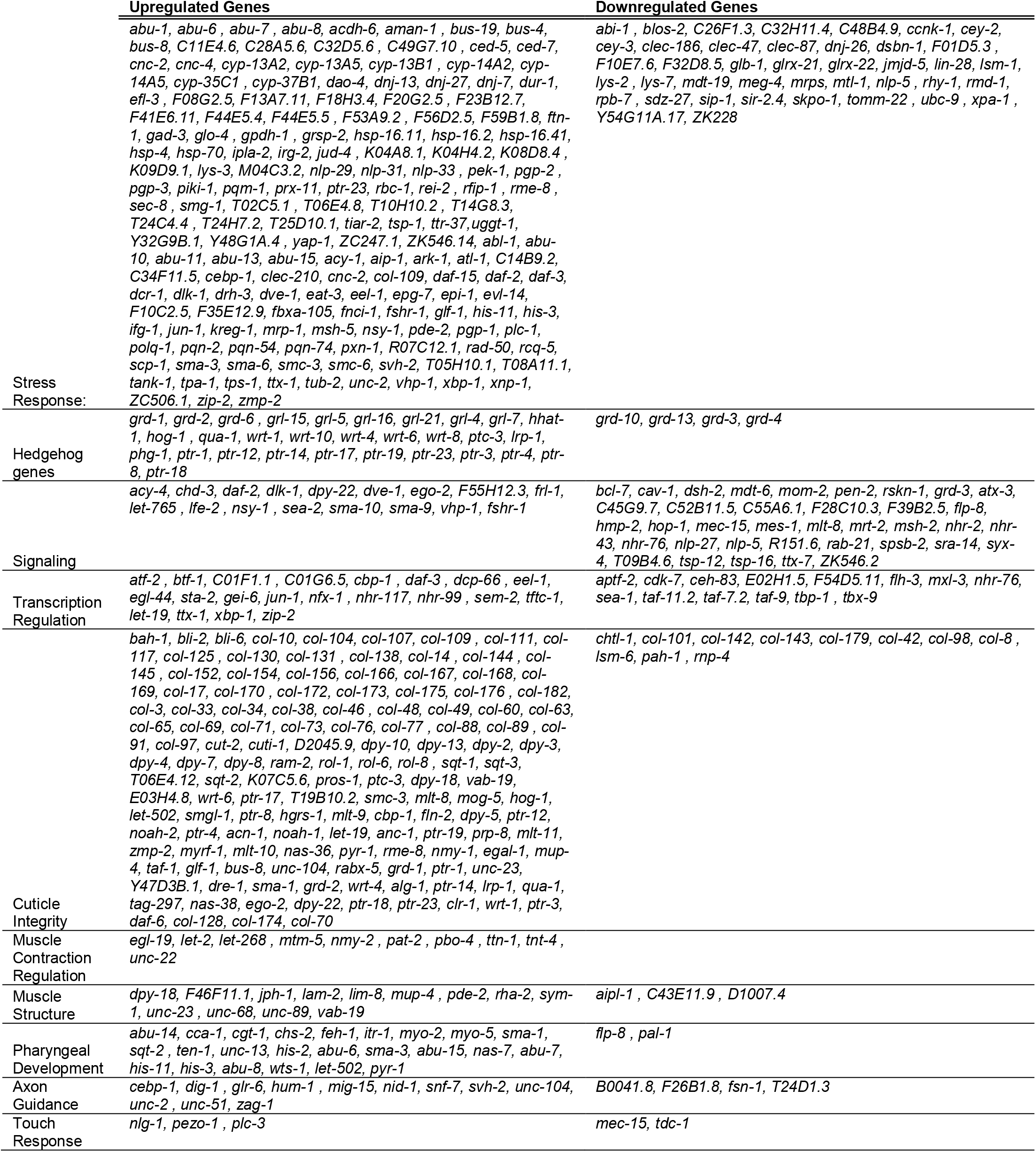
Representative gene clusters that were altered as a result of mechanical strain application in wild-type *C. elegans*.

**Table S2:**
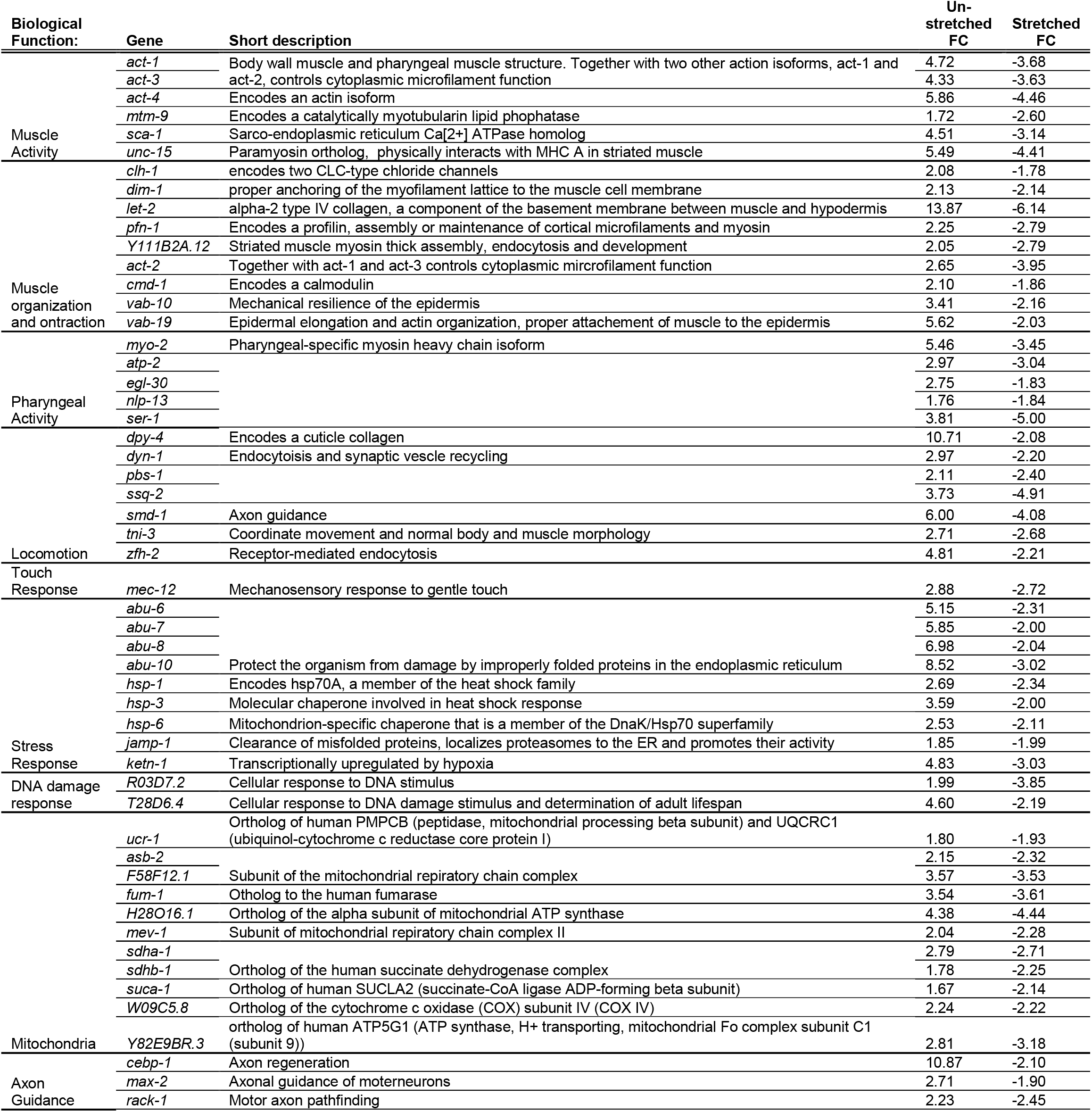
Key genes involved in the compensatory response of L535P animals to mechanical strain application. Shown are genes that were downregulated in un-strained animals and specifically upregulated in response to animals being stretched.

**Table S3:**
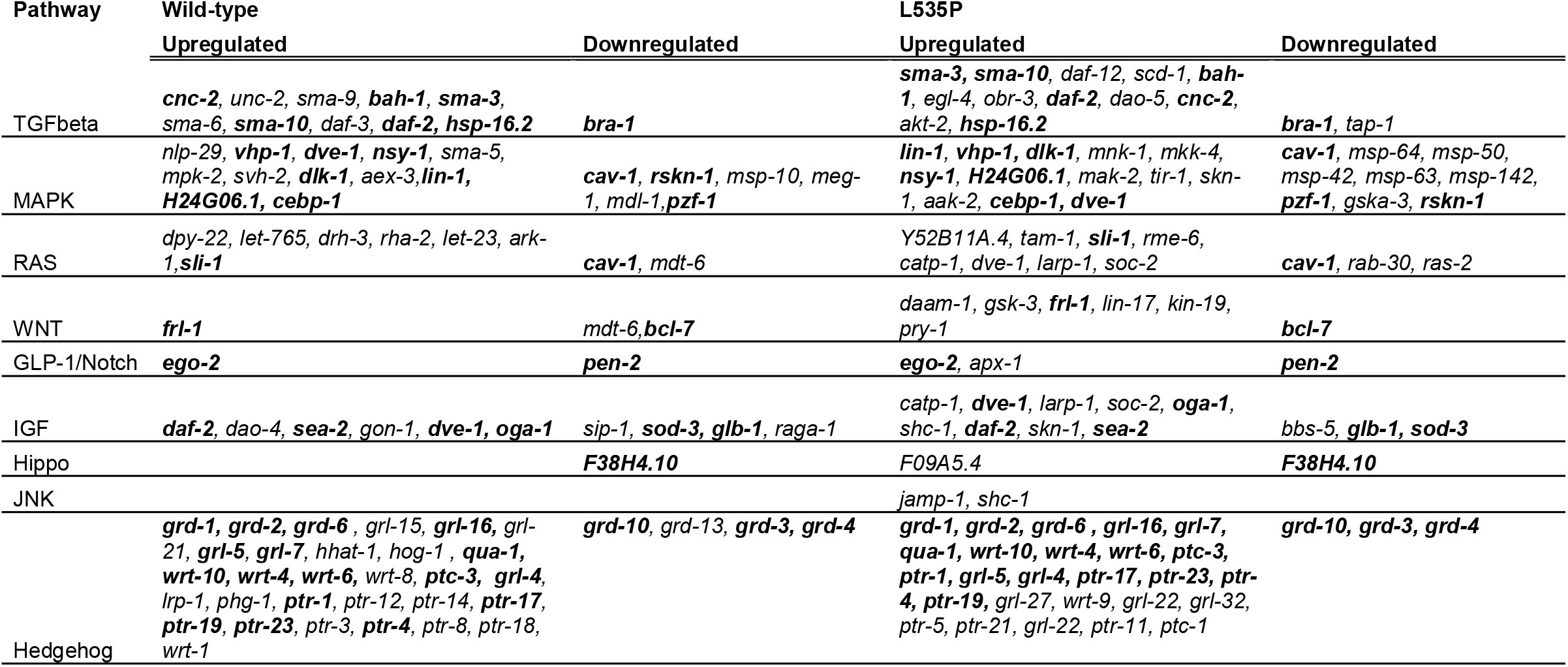
Genes involved in molecular pathways altered by external strain application in both wild-type and L535P expressing animals. Common genes are indicated in bold.

**Table S4:** Percent mortality in mechanically stretched worms

| Name | Age in days at 50 % mortality |
| --- | --- |
| Un-stretched | 14 |
| 5 minute stretch | 15 |
| 15 minute stretch | 12 |
| 30 minute stretch | 4 |

**(B)** Fisher's exact test
| Condition | P value at- |  |  |  |
| --- | --- | --- | --- | --- |
|  | 25% mortality | 50% mortality | 75% mortality | 95% mortality |
| Un-stretched vs. 5 minute stretch | 0.0032 | 0.0002 | 0.5892 | 0.6842 |
| Un-stretched vs. 15 minute stretch | 1.9 e-8 | 0.0202 | 0.0623 | 0.0289 |
| Un-stretched vs. 30 minute stretch | 1.5 e-12 | 1.2 e-12 | 1.1 e-12 | 6.2 e-13 |
| 5 minute stretch vs. Un-stretched | 0.0032 | 0.0002 | 0.5892 | 0.6842 |
| 5 minute stretch vs. 15 minute stretch | 7 e-6 | 0.0004 | 0.0294 | 0.0162 |
| 5 minute stretch vs. 30 minute stretch | 4.9 e-13 | 4.9 e-13 | 1.5 e-12 | 1.1 e-12 |
| 15 minute stretch vs. Un-stretched | 1.9 e-8 | 0.0202 | 0.0623 | 0.0289 |
| 15 minute stretch vs. 5 minute stretch | 7 e-6 | 0.0004 | 0.0294 | 0.0162 |
| 15 minute stretch vs. 30 minute stretch | 9.4 e-13 | 9.4 e-13 | 8.4 e-13 | 6.4 e-9 |
| 30 minute stretch vs. Un-stretched | 1.5 e-12 | 1.2 e-12 | 1.1 e-12 | 6.2 e-13 |
| 30 minute stretch vs. 5 minute stretch | 4.9 e-13 | 4.9 e-13 | 1.5 e-12 | 1.1 e-12 |
| 30 minute stretch vs. 15 minute stretch | 9.4 e-13 | 9.4 e-13 | 8.4 e-13 | 6.4 e-9 |

***mec-3* (RNAi):****(A) Mean Lifespan**
| Name | Age in days at 25 % mortality |
| --- | --- |
| WT *EV* un-stretched | 12 |
| WT *mec-3* (RNAi) un-stretched | 12 |
| WT *EV* stretched | 10 |
| WT *mec-3* (RNAi) stretched | 10 |

**(B) Fisher's exact test**
| Condition | P value at- |  |  |  |
| --- | --- | --- | --- | --- |
|  | 25% mortality | 50% mortality | 75% mortality | 95% mortality |
| *EV* un-stretched vs. *EV* stretched | 0.0001 | 0.0324 | 0.6408 | 0.4996 |
| *EV* un-stretched vs. *mec-3* (RNAi) un-stretched | 0.1008 | 0.316 | 0.316 | 0.316 |
| *EV* un-stretched vs. *mec-3* (RNAi) stretched | 0.0313 | 0.0166 | 0.1276 | 0.1276 |
| *EV* stretched vs. *EV* un-stretched | 0.0001 | 0.0324 | 0.6408 | 0.4996 |
| *EV* stretched vs. *mec-3* (RNAi) un-stretched | 0.224 | 0.433 | 0.4208 | 0.4358 |
| *EV* stretched vs. *mec-3* (RNAi) stretched | 0.449 | 0.3201 | 0.16 | 0.1739 |
| *mec-3* (RNAi) un-stretched vs. *EV* un-stretched | 0.1008 | 0.316 | 0.316 | 0.316 |
| *mec-3* (RNAi) un-stretched vs. *EV* stretched | 0.224 | 0.433 | 0.4208 | 0.4358 |
| *mec-3* (RNAi) un-stretched vs. *mec-3* (RNAi) stretched | 0.8239 | 0.167 | 0.6584 | 0.6584 |
| *mec-3* (RNAi) stretched vs. *EV* un-stretched | 0.0313 | 0.0166 | 0.1276 | 0.1276 |
| *mec-3* (RNAi) stretched vs. *EV* stretched | 0.224 | 0.433 | 0.4208 | 0.4358 |
| *mec-3* (RNAi) stretched vs. *mec-3* (RNAi) un-stretched | 0.8239 | 0.167 | 0.6584 | 0.6584 |

**(A) Mean Lifespan**
| Name | Age in days at 25 % mortality |
| --- | --- |
| WT *EV* un-stretched | 12 |
| WT *mec-15* (RNAi) un-stretched | 12 |
| WT *EV* stretched | 10 |
| WT *mec-15* (RNAi) stretched | 10 |

**(B) Fisher's exact test**
| Condition | P value at- |  |  |  |
| --- | --- | --- | --- | --- |
|  | 25% mortality | 50% mortality | 75% mortality | 95% mortality |
| *EV* un-stretched vs. *EV* stretched | 0.0001 | 0.0324 | 0.6408 | 0.4996 |
| *EV* un-stretched vs. *mec-15* (RNAi) un-stretched | 0.3808 | 0.2724 | 1 | 0.8862 |
| *EV* un-stretched vs. *mec-15* (RNAi) stretched | 0.0095 | 0.0172 | 0.0172 | 0.0059 |
| *EV* stretched vs. *EV* un-stretched | 0.0001 | 0.0324 | 0.6408 | 0.4996 |
| *EV* stretched vs. *mec-15* (RNAi) un-stretched | 0.0109 | 0.6013 | 0.5579 | 0.737 |
| *EV* stretched vs. *mec-15* (RNAi) stretched | 0.2115 | 0.2082 | 0.116 | 0.0049 |
| *mec-15* (RNAi) un-stretched vs. *EV* un-stretched | 0.3808 | 0.2724 | 1 | 0.8862 |
| *mec-15* (RNAi) un-stretched vs. *EV* stretched | 0.0109 | 0.6013 | 0.5579 | 0.737 |
| *mec-15* (RNAi) un-stretched vs. *mec-15* (RNAi) stretched | 0.0213 | 0.1748 | 0.011 | 0.0099 |
| *mec-15* (RNAi) stretched vs. *EV* un-stretched | 0.0095 | 0.172 | 0.172 | 0.0059 |
| *mec-15* (RNAi) stretched vs. *EV* stretched | 0.2115 | 0.2082 | 0.116 | 0.0049 |
| *mec-15* (RNAi) stretched vs. *mec-15* (RNAi) un-stretched | 0.0213 | 0.1748 | 0.011 | 0.0099 |

**(A) Mean Lifespan**
| Name | Age in days at 25 % mortality |
| --- | --- |
| WT *EV* un-stretched | 12 |
| WT *pezo-1* (RNAi) un-stretched | - |
| WT *EV* stretched | 10 |
| WT *pezo-1* (RNAi) stretched | - |

**(B) Fisher's exact test**
| Condition | P value at- |  |  |  |
| --- | --- | --- | --- | --- |
|  | 25% mortality | 50% mortality | 75% mortality | 95% mortality |
| *EV* un-stretched vs. *EV* stretched | 0.0001 | 0.0324 | 0.6408 | 0.4996 |
| *EV* un-stretched vs. *pezo-1* (RNAi) un-stretched | 0.4551 | 0.3006 | 0.0001 | 0.0001 |
| *EV* un-stretched vs. *pezo-1* (RNAi) stretched | 3.9e-6 | 5.2e-6 | 0.0136 | 0.0858 |
| *EV* stretched vs. *EV* un-stretched | 0.0001 | 0.0324 | 0.6408 | 0.4996 |
| *EV* stretched vs. *pezo-1* (RNAi) un-stretched | 5.4e-7 | 0.0006 | 6.3e-8 | 9e-10 |
| *EV* stretched vs. *pezo-1* (RNAi) stretched | 0.0278 | 0.3063 | 0.3675 | 0.001 |
| *pezo-1* (RNAi) un-stretched vs. *EV* un-stretched | 0.4551 | 0.3006 | 0.0001 | 0.0001 |
| *pezo-1* (RNAi) un-stretched vs. *EV* stretched | 5.4e-7 | 0.0006 | 6.3e-8 | 9e-10 |
| *pezo-1* (RNAi) un-stretched vs. *pezo-1* (RNAi) stretched | 0.0009 | 1.4e-7 | 0.0002 | 0.0021 |
| *pezo-13* (RNAi) stretched vs. *EV* un-stretched | 3.9e-6 | 5.2e-6 | 0.0136 | 0.0858 |
| *pezo-1* (RNAi) stretched vs. *EV* stretched | 0.0278 | 0.3063 | 0.3675 | 0.001 |
| *pezo-1* (RNAi) stretched vs. *pezo-1* (RNAi) un-stretched | 0.0009 | 1.4e-7 | 0.0002 | 0.0021 |

**(A) Mean Lifespan**
| Name | Age in days at 50 % mortality |
| --- | --- |
| WT *EV* un-stretched | - |
| WT *ptc-3* (RNAi) un-stretched | - |
| WT *EV* stretched | 5 |
| WT *ptc-3* (RNAi) stretched | 3 |

**(B) Fisher's exact test**
| Condition | P value at- |  |  |  |
| --- | --- | --- | --- | --- |
|  | 25% mortality | 50% mortality | 75% mortality | 95% mortality |
| *EV* un-stretched vs. *EV* stretched | 7.2 e-13 | 7.2 e-13 | 7.2 e-13 | 7.2 e-13 |
| *EV* un-stretched vs. *ptc-3* (RNAi) un-stretched | 5.2 e-13 | 1.3 e-12 | 1.9 e-12 | 1.1 e-12 |
| *EV* un-stretched vs. *ptc-3* (RNAi) stretched | 1.6 e-12 | 1.6 e-12 | 9.2 e-13 | 6.6 e-13 |
| *EV* stretched vs. *EV* un-stretched | 7.2 e-13 | 7.2 e-13 | 7.2 e-13 | 7.2 e-13 |
| *EV* stretched vs. *ptc-3* (RNAi) un-stretched | 1.3 e-12 | 1.3 e-12 | 0.0064 | 1.6 e-12 |
| *EV* stretched vs. *ptc-3* (RNAi) stretched | 1.6 e-12 | 1.6 e-12 | 1.6 e-12 | 1.8 e-12 |
| *ptc-3* (RNAi) un-stretched vs. *EV* un-stretched | 5.2 e-13 | 1.3 e-12 | 1.9 e-12 | 1.1 e-12 |
| *ptc-3* (RNAi) un-stretched vs. *EV* stretched | 1.3 e-12 | 1.3 e-12 | 0.0064 | 1.6 e-12 |
| *ptc-3* (RNAi) un-stretched vs. *ptc-3* (RNAi) stretched | 1.6 e-12 | 1.6 e-12 | 1.3 e-12 | 6.5 e-10 |
| *ptc-3* (RNAi) stretched vs. *EV* un-stretched | 1.6 e-12 | 1.6 e-12 | 9.2 e-13 | 6.6 e-13 |
| *ptc-3* (RNAi) stretched vs. *EV* stretched | 1.6 e-12 | 1.6 e-12 | 1.6 e-12 | 1.8 e-12 |
| *ptc-3* (RNAi) stretched vs. *ptc-3* (RNAi) un-stretched | 1.6 e-12 | 1.6 e-12 | 1.3 e-12 | 6.5 e-10 |

